# A database for ITS2 sequences from nematodes

**DOI:** 10.1101/689695

**Authors:** Matthew L. Workentine, Rebecca Chen, Shawna Zhu, Stefan Gavriliuc, Nicolette Shaw, Jill de Rijke, Elizabeth M. Redman, Russell W. Avarmenko, Janneke Wit, Jocelyn Poissant, John S. Gilleard

## Abstract

Marker gene surveys have a wide variety of applications in species identification, population genetics, and molecular epidemiology. As these methods expand to new types of organisms and additional markers beyond 16S and 18S rRNA genes, comprehensive databases are a critical requirement for proper analysis of these data. Here we present an ITS2 rDNA database for maker gene surveys of both free-living and parasitic nematode populations and the software used to build the database. This is an important resource for researchers working on nematodes and also provides a tool to create ITS2 databases for any given taxonomy.

**Availability and Implementation:** The database is available as an interactive web app at https://cooperia.chgi.ucalgary.ca/Nematode_ITS2/. The full database can also be downloaded from zenodo https://doi.org/10.5281/zenodo.3235802, and the open source software used to create the database, markerDB is available at https://github.com/ucvm/markerDB/releases/latest.

## Introduction

The ITS2 rDNA locus has been widely used as marker for species identification in both free-living and parasitic nematodes for many years (refs). Nematodes, as with other invertebrate groups, often exist in large and complex communities. Consequently, deep amplicon sequencing approaches have a potentially powerful role for the investigation nematode communities similar to the use of bacterial 16S rDNA amplicon sequencing in microbiome studies. For example, the ITS2 rDNA locus has recently been used for “nemabiome” sequencing of parasitic nematode communities inhabiting the gastrointestinal tract of cattle (Avramenko *et al.*, 2017). In that case, reliable species identification was achieved using a small bespoke, curated ITS2 rDNA database of the major relevant cattle gastrointestinal nematode species. However, the wider and more versatile application of deep amplicon sequencing approaches to nematode research will requires a more comprehensive, and regularly updated, ITS2 rDNA database equivalent to that available for studying fungi (Nilsson *et al.*, 2019). In this paper, we describe the development of a nematode ITS2 rDNA database.

## Development

The nemtatode ITS2 database was constructed using markerDB, which we have provided as an open-source tool to quickly and reliably construct an ITS2 database for any NCBI taxonomic level. This tool is made available to facilitate reproducability and transparency, and to provide users with the option to construct their own databases. markerDB is implemented in the R programming language and run as a Snakemake pipeline. A brief description of the pipeline follows.

Potential ITS2 sequences are retrieved from NCBI using the rentrez R package (Winter, 2017) based on a text search that will find ITS2 annotated sequences that are limited to the provided taxonomy. The full taxonomy of the downloaded sequences is obtained with the taxize R package (Chamberlain *et al.*, 2019; Scott Chamberlain and Eduard Szocs, 2013). Only taxonomies that are complete with all ranks (Superkingdom, Kingdom, Phylum, Class, Order, Family, Genus, Species) are retained. Additionally, taxonomies with incomplete species names, which contain numbers or ‘sp.’ are removed.

Many of the ITS2 annotated sequences also contain the partial or full upstream and downstream 5.8S and 28S genes and so trimming to the ITS2 region is required. However, no good sequence models exist that capture a wide range of diversity due to the divergence of this region. For this step Infernal, (specifically cmscan) (Nawrocki and Eddy, 2013) is used to identify the 5.8S and eukaryotic LSU genes. The covariance models used to create the nematode database and also provided with markerDB, were taken from Rfam (Kalvari *et al.*, 2018). If a hit to the 5.8S is identified (parital hits allowed) this region and everything upstream is trimmed off. This is repeated downstream for any 28S hits. Any retrieved sequences that don’t have hits to either rRNA gene are assumed to be soley ITS2 and are retained in the database. This option can be changed when running the pipeline. Finally, sequences too long or too short (700 bp and 100 bp, respectivaly, as set in the configuration) are discarded. The final sequence set contains a fair bit of redundancy and so a non-redundant version of the database with unique sequences only is returned. If an alignment is required an option to align the sequences using MAFFT (Katoh *et al.*, 2002) is also provided but it should be noted that aligning ITS2 sequences from diverse organisms is difficult due to the heterogeneity present and in general we recommend taxonomy assignment methods that do not depend on alignments, particularly for databases covering a large taxonomic range.

The output of markerDB is a fasta file with the final sequences and a coresponding tab-delimited text file with the taxonomy, linked by Genbank accession number. The pipeline also provides function to write out the database in formats used with popular taxonomy assignment methods including dada2 (Callahan *et al.*, 2016), the rRDP Bioconductor package (Hahsler and Nagar, 2019), mothur (Schloss *et al.*, 2009), and IDTAXA (Murali *et al.*, 2018). For example the IDTAXA output files can be used with our recommended nemabiome analysis workflow (www.nemabiome.ca). A simple shiny app is also provided that allows users to work with the database interactively, filtering taxonomic groups as needed and downloading the filtered data in any of the above formats.

## Summary

The database (version 1.0.0 at the time of writing) currently containts 8630 non-redunant sequences with a median length of 263 bp and standard deviation of 97 bp. There are 1429 species and 325 genera and across the taxonomic ranks we were able to obtain good quality, non-redundant sequences for approximately 30% of the taxa in the NCBI database in the Nematoda phylum (see Figure 1).

**Figure 1:**
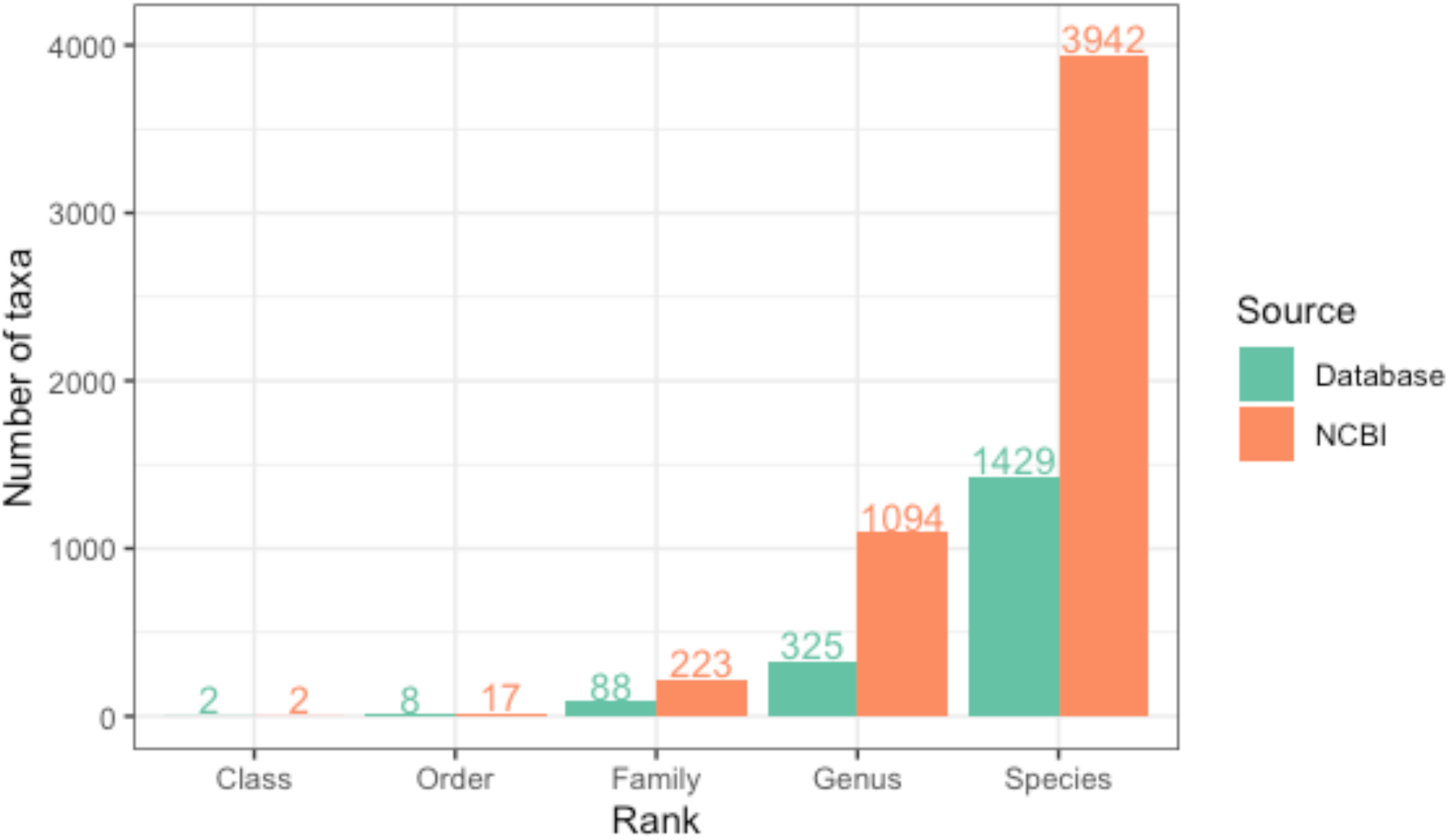
Number of taxa in the database compared to the total in the NCBI taxonomy database

We have also provided a simple web app, which allows users to search and filter the database and create versions customized to their research area of interest. Futher, the database will be updated every 3-6 months, feasible due the automated and reproducibility of the database construction using markerDB. Rapid updates allow researchers to generate analysis that reflect the most current sequences in Genbank.

In conclusion, we provide a database of nematode ITS2 sequences that greatly expands the range of sequences suitable to study both parasitic and free-living nematode communities allowing a broader selection of hosts and environments to be studied. We have also provided open source software to easily and reproduceably build ITS2 databases for any taxonomy of interest.

